# GiRAFR improves gRNA detection and annotation in single cell CRISPR screens

**DOI:** 10.1101/2022.10.24.513352

**Authors:** Qian Yu, Paulien Van Minsel, Eva Galle, Bernard Thienpont

## Abstract

Novel single cell RNA-seq analysis combined with CRISPR screens enables the high-throughput characterization of transcriptional changes caused by genetic perturbations. Dedicated software to annotate CRISPR guide RNA (gRNA) libraries and associate them with single cell transcriptomes are however lacking. Here, we generated a CRISPR droplet sequencing dataset. We demonstrate that the current default tool fails to detect mutant gRNAs. We therefore developed GiRAFR, a pysam-based software tool to characterize intact and mutant gRNAs. We show that mutant gRNAs are dysfunctional, and failure to detect and annotate them leads to an inflated estimate of the number of untransformed cells as well as an underestimated multiplet frequency. These findings are mirrored in publicly available datasets, where we find that up to 34 % of cells are transduced with a mutant gRNA. Applying GiRAFR hence stands to improve the annotation and quality of single cell CRISPR screens.

## Introduction

The advent of single cell omics technologies is revolutionizing the cataloguing of cell types and states in development, physiology, and disease, with the Human Cell Atlas epitomizing many of these efforts. While these studies are mostly descriptive and correlative, recent method developments now also enable functional analysis at scale, with the impact of a multitude of gene perturbations for the first time assessed in parallel on a transcriptome-wide scale. Examples include Perturb-seq, Direct-Capture Perturb-seq, CROP-seq, ECCITE-seq and others (1–5). Screens initially encompassed dozens of parallel perturbations, but more recent iterations profiled numbers that are several orders of magnitude higher, extending to the whole transcriptome (6–8). These screens mostly rely on CRISPR-based systems where guide RNAs (gRNA) are used to target Cas9 to loci of interest for gene knockout, inactivation or overexpression. The concomitant read-out of a single cell’s mRNA and gRNA expression allows for a high-through characterization of genetic perturbation phenotypes. Current data analyses pipelines are however poorly catered for these types of analyses, and also quality control metrics for such experiments have yet to be established.

Here, we developed GiRAFR (Guide RNA Anomalies and Functionality Revealed), a tool based on pysam (9) to perform quality control of single cell CRISPR screens, and to assign gRNAs to cells in a sensitive, mutation-aware manner. This analysis enables profiling of gRNA sequence variations, as well as allowing to pinpoint the sources of this variation, such as induced during library preparation or during virus preparation. In a separate analysis mode, this tool also detects CRISPR-cas9-induced DNA editing in transcriptome data. By applying GiRAFR to 1 new and 11 publicly available datasets (26 experiments), we show that cells are often inaccurately annotated, and we propose minimal quality metrics for single cell gRNA sequencing libraries. Together, GiRAFR represents a toolbox to analyze single cell CRISPR screens in a more accurate, reliable, variance-aware and efficient manner.

## Results

We transformed A549 cells expressing a tamoxifen-inducible Cas9 with a lentiviral pool for expression of 120 gRNAs (Table S1) at low multiplicity of infection. After stringent selection using puromycin and 2 days of Cas9 induction, cells were allowed to grow for another 5 days. Next, 5,744 cells were analyzed by CROP-seq as described. Downstream analysis confirmed this experiment to be successful, as cells carrying specific gRNAs showed downregulated expression of gRNA target genes [Figure 1a]. To further validate functionality, we searched for Cas9-induced indels at gRNA target regions (see methods). Also this confirmed that our experiment performed as anticipated, since indels were readily identified in highly expressed genes when the gRNA target region was recovered in the transcriptome library [Figure 1b]. Surprisingly however, no gRNA was detected in 2,317 cells using the standard gRNA detection and annotation pipeline, Cell Ranger (10) [Figure S1a]. Single cell CRISPR screens are contingent on successfully and accurately associating a cell’s transcriptome with the gRNA it expresses. Similar to what we observed here however, in every study published thus far, no gRNA is detected in a subset of cells (6–8,11,12). This is typically attributed to a lack of complete selection for gRNA-transformed cells, leading to inclusion of non-transformed cells in the analysis, or to an insufficiently high gRNA expression or an insufficient sequencing depth, implying that a gRNA is expressed but not detected. In our experiment however, sequencing depth was high, with a saturation estimated at 98.9 %, and also gRNA expression in gRNA-positive cells was high, with on average 53 unique molecular identifiers (UMIs) per gRNA per cell [Figure S1b]. Surprisingly, also gRNA-negative cells showed expression of the puromycin resistance cassette we used as selection marker, suggesting that these cells were successfully transformed and selected [Figure 1c]. Closer inspection of reads mapping to the gRNA-expressing plasmid sequence moreover revealed that also gRNA-derived reads are mapped to this region. These however show imperfect mapping, suggesting the presence of gRNA mutations [Figure 1d].

**Figure 1.**
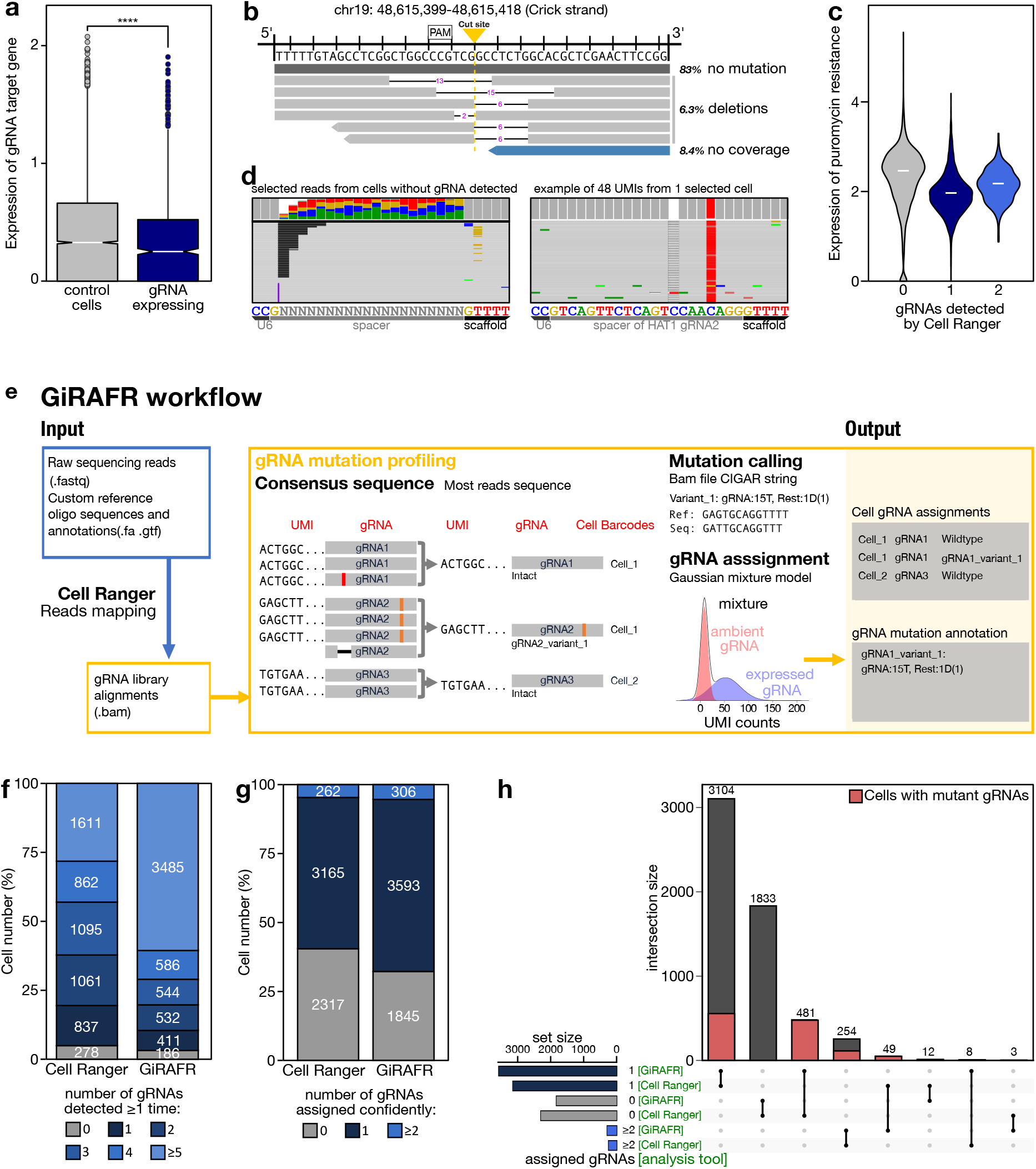
Development of GiRAFR. **a**. Expression (log-transformed read counts) of gRNA target genes, normalized to their expression in cells with non-targeting gRNAs (control cells). Shown is the aggregate expression of the 11 target genes for which the corresponding gRNA is detected in at least 70 cells, and that show expression in at least 50 % of all control cells. ****: P < 0.0001 by 2-sided t-test. **b**. 51 bp window of *RPL18*, showing the predicted cut site of gRNA 1 (targets the Crick strand). Nine cells expressing this gRNA show reads coverage in this detection window. 6 of 95 reads (6.3 %) with a unique UMI show a deletion across the cut site (3-4bp upstream of PAM), 83% show no mutations, and 8.4% have no coverage over the cut site. Note that transcripts with out-of-frame indels are likely subject to nonsense-mediated decay. These frequencies therefore likely underestimate the genome editing efficiency. **c**. Expression of the Puromycin resistance cassette, after mapping the scRNA-seq library to a reference augmented to include the Puromycin resistance cassette. Shown is the log-normalized expression in cells where 0, 1 or 2 gRNAs were detected by Cell Ranger feature barcoding analysis. **d**. Reads with unique UMIs showing partial mapping to the 20 bp spacer region, using either Ns as a reference (left), or from 1 cell expressing a variant *HAT1* gRNA using this gRNA as a reference. **e**. Schematic workflow of the GiRAFR pipeline. See methods for additional details. **f-g**. Number of cells with gRNA spacer assigned by Cell Ranger and GiRAFR (f) in the raw count matrix and (g) after application of the Gaussian Mixture model represented panel e. **h**. Comparison of gRNA assignment to cells between Cell Ranger and GiRAFR (20). The red box indicates cells containing 1 or more mutant gRNAs.

Detection and annotation of such gRNA mutations is of importance for several reasons. Firstly, for controlling the quality of the experimental steps leading up to gRNA detection such as gRNA oligonucleotide synthesis and cloning. Secondly, for understanding the actual performance of cellular transformation and selection. And finally, and perhaps most importantly, because single cell CRISPR screens often rely on gRNAs for an accurate discrimination between single cells, having one gRNA, and so-called multiplets, where two or more cells are inadvertently captured together causing them to share barcodes. Such multiplets as well as double-transformed cells are identified as having more than one gRNA and are typically discarded for analysis. Indeed, if such a multiplet is incorrectly labelled as a single cell, the corresponding transcriptome no longer reflects the impact of the CRISPR perturbation. The current state-of-the-art pipeline for assigning gRNAs to cells is the feature barcoding analysis implemented in Cell Ranger. It is however unable to identify mutant gRNAs that are over 1 Hamming distance away from the designed gRNA sequence. To remedy this, we developed GiRAFR, a tool to identify mutations in the gRNA expression library and assign intact and mutant gRNAs to cells in single cell screens. It includes multiple controllable filtering and model fitting parameters to establish accurate spacer calling and provides information on both intact and mutant gRNAs [Figure 1e]. Specifically, it first calls a consensus gRNA sequence for each UMI. Next, it generates a count matrix per detected gRNA, whilst removing UMIs with fewer than 2 reads to avoid including consensus gRNAs reflecting sequencing errors. Next, by aligning the consensus gRNA sequences to the predefined gRNA library, mutant and intact gRNAs can be identified and mutations annotated (see methods). Here, we hence assembled a library of in total 1,997 different gRNAs (113 wild-type and 1,884 mutant). Once this listing of consensus gRNA sequences was constructed, gRNAs were assigned to cells using the corresponding cell barcode. In each cell, we detected between 0 to over 40 different gRNAs, versus 0 to 22 gRNAs for Cell Ranger [Figure 1f, Figure S1c]. In each cell we detected on average 43 UMIs from 7 different gRNAs, either intact or mutant. After filtering out UMIs with only 1 read, some of these remaining gRNAs are supported by multiple different UMIs in a cell, others have only 1 UMI and thus represent single gRNA molecules. These can originate from so-called ambient RNA, released from cells lysed prior to encapsulation and inadvertently captured together with another unrelated cell, but also from errors in gRNA production in the cell or its conversion to DNA during library preparation. The latter scenario represents a minority of all instances, as only in 1,100 of 5,359 cells, both the mutant and intact version of the same gRNA was encountered. In most of these instances (93 %), the mutant version was supported by only 1 UMI.

Irrespective of this, deciding which gRNAs are truly expressed in a cell and which represent artifacts, is a common problem in single cell CRISPR screen analysis. In GiRAFR, we therefore implemented the two-components Gaussian mixture model also used by Cell Ranger, to determine dynamic UMI thresholds per gRNA per cell (Figure 1e). In this implementation, we included both mutant and intact gRNA UMI counts to model the distribution, implying that different thresholds may be proposed in GiRAFR *versus* Cell Ranger. Irrespective of this, GiRAFR detects more cells with a gRNA (Figure 1g). Indeed, when assigning gRNAs to cells using GiRAFR, 481 of the 2,317 cells where no gRNA was previously found, appeared transformed with a mutant gRNA, and 49 cells where previously only one gRNA was found were in fact multiplets (Figure 1h). In total 1,187 of all 5,744 cells included in the analysis (20,7 %) contain a mutant gRNA.

A key question is if gRNA mutations affect their function. The most frequent mutation is deletion of a thymidine in the TTTT tetramer of the gRNA lower stem structure, but mutations occur both in the gRNA promoter, spacer and scaffold, and could thus compromise gRNA expression, DNA target recognition and Cas9 binding, respectively (13) (Figure 2a). To evaluate the functionality of both intact and mutant gRNAs, we compared the transcript level of all 11 sufficiently expressed gRNA target genes between non-perturbed cells and cells perturbed with a mutant or intact gRNA (Figure S2, Figure 2b). While intact gRNAs reduced target gene expression by on average 19.8 % (P = 2.3 × 10^−10^), gRNAs with a mutant scaffold induced a 12.1% reduction in expression (P = 0.0498) and gRNAs with a mutant spacer reduced expression by only 7.6% (not significant). Mutant gRNAs thus appear to be dysfunctional.

**Figure 2.**
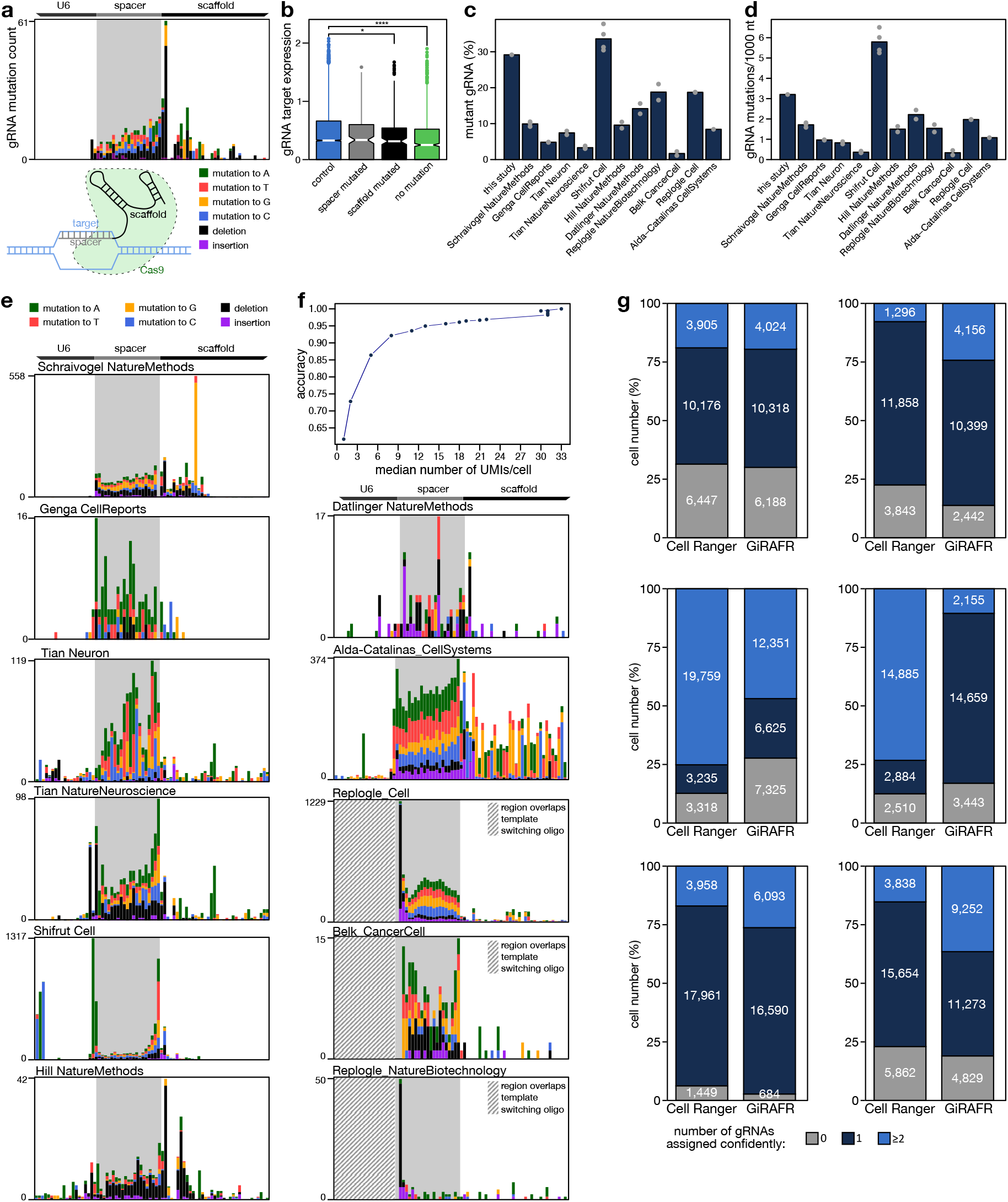
Application of GiRAFR to an extended dataset. **a**. Mutation frequency of gRNAs associated with cells after the Gaussian Mixture model filtering. The cartoon below illustrates the annotation of spacer and scaffold in the gRNA. **b**. Expression (log-transformed read counts) of gRNA target genes, normalized to their expression in cells with non-targeting gRNAs (control cells). Shown is the aggregate expression of the 11 target genes for which the corresponding gRNA is detected in at least 70 cells, and that show expression in at least 50 % of all control cells. *, ****: P < 0.05, 0.0001 by a 2-sided t-test. **c**. Fraction of cells with a gRNA showing a mutation, out of all cells. Only gRNAs associated with cells after the Gaussian Mixture model filtering were considered. **d**. Frequency of mutations in gRNAs associated with cells after the Gaussian Mixture model filtering. The mutation frequency is shown per cell and per 1,000 nucleotides, to subtract differences in gRNA sequencing read length. **e**. gRNA mutation patterns detected in publicly available datasets, as in panel a. Shown is the aggregate across all experiments analyzed per dataset. Note that some gRNA libraries encompass only a fraction of the U6 promoter shown. Individual experiments are shown in Figure S2. Striped boxes indicate the position of template switching oligonucleotides, where mutations on position −2, −1 and 0 before the start of the spacer were also removed. **f**. gRNA assignment accuracy as a function of the median number of UMIs per cell, as estimated by downsampling the CROP-seq data generated in the current study. **g**. Comparison of the number of different gRNAs assigned to a cell by Cell Ranger and GiRAFR. Shown are (left and right) data from references 12 and 2 (top), 22 and 8 (middle), and 21 and 14 (bottom).

As described higher, GiRAFR is able to detect both mutant and intact gRNAs, and can accurately assign them to cells. This explains why a subset of cells appeared to be untransformed with gRNA-expressing plasmid in our experiment, and highlights that some cells annotated as singlets are in fact multiplets. To assess if such mutant gRNAs also affect other, independent experiments, we applied GiRAFR with default configuration to 26 single cell CRISPR experiments from 11 studies (Table S2). While the gRNA library in each study was constructed with a different strategy, resultant sequences usually contain both a region before the spacer (i.e. the end of U6 promoter or the TSO), the spacer region itself that contains the unique gRNA sequence (~ G + 20bp), and the remainder, representing gRNA scaffold and/or capture sequence. The percentage of mutant gRNA molecules varies across different studies from 2 % to 34 %, with different samples from the same study showing concordant results (Figure 2c). Mutation spectra were different across studies, with mutation frequencies ranging from 0.2 to 5.8 mutations per 1,000 nucleotides per gRNA profiled (Figure 2d). Frequencies were higher in the spacer region and in regions immediately flanking it, corresponding to the sequences that are typically synthesized as pooled oligonucleotides for generating gRNA virus libraries (Figure 2e). Mutations in addition frequently occurred in the beginning and at the end of the spacer region, sites that serve as handles for cloning these oligonucleotides into the gRNA expression plasmid prior to virus production.

To assess the impact of these gRNA mutations on gRNA assignment and on identification of cells transformed with a single gRNA, we compared results from GiRAFR with Cell Ranger feature barcoding analysis. When no separate gRNA enrichment library was available, we provided Cell Ranger with the scRNA-seq library as CRISPR Guide Capture Library. Notably, the gRNA library sequencing saturation between these studies differed dramatically, ranging from 20 to 99 %. Also the average number of gRNA molecules detected per cell was highly variable, ranging from 1 to over 100 UMIs (Figure S2b). We first assessed the impact of this variation on the anticipated outcome, by down-sampling our own CROP-seq experiment. This analysis revealed that, at a sequencing saturation of 34, 59 and 92 %, an average of 5, 16 and 31 unique gRNA molecules were recovered per cell, leading to a correct annotation of cells with their gRNA at an estimated 80%, 95% and 98% of samples (Figure 2f). In 13 of 26 experiments we analyzed, on average over 16 unique gRNA molecules were recovered per cell, suggesting that over 95 % of cells are correctly assigned. We focused our analyses on these 13 experiments (6 studies); in the 13 other experiments, many cells were excluded from analysis by GiRAFR as a gRNA cannot be confidently assigned to cells. In each of the 13 high-depth experiments however, GiRAFR identified more cells as containing a mutant gRNA (Figure S3). Note that not each of the studies we analyzed implemented gRNA analysis using the same Gaussian Mixture model to define UMI thresholds. Indeed, they often use *ad hoc*, fixed thresholds, rendering a head-to-head comparison between each of these models and GiRAFR unfeasible. To enable a head-to-head comparison, we therefore reanalyzed each of these datasets using Cell Ranger as well as GiRAFR. This revealed that Cell Ranger labelled between 6 and 39 % of all cells as containing no gRNA, but 0.5 to 55 % of these in fact represent cells transfected with mutant gRNAs as per the GiRAFR analysis. Likewise, cells labelled as having a single gRNA by Cell Ranger were often labelled as multiplets by GiRAFR, with for example about 1 in 3 cells profiled by Replogle *et al*. (14) being mislabeled by Cell Ranger (Figure 2g). These analyses confirm that GiRAFR outperforms Cell Ranger for gRNA library analysis and for assigning gRNAs to cells.

## Discussion

Single cell CRISPR screening is increasingly being used to assess the impact of gene perturbations on cellular transcriptomes. Here, we developed a method to identify and annotate the gRNAs detected in these assays more accurately. Our approach is useful in several respects. First and foremost, if mutant gRNAs cannot be detected, cells with a single gRNA cannot be properly distinguished from those with multiple gRNAs. Discriminating both sets of cells is important to avoid including cells that are inadvertently captured together and labelled with the same barcode. In that scenario, the transcriptome no longer represents the associated perturbation. We demonstrate that up to 35 % of all cells profiled in earlier studies that are labelled as singlets by Cell Ranger are in fact cell multiplets, i.e. multiple cells, or single cells transformed with 2 or more gRNAs. Note that these studies do not always rely on the same Gaussian Mixture model to define gRNA UMI thresholds, but often use *ad hoc* customized pipelines or fixed thresholds, rendering a head-to-head comparison between each of these models and GiRAFR unfeasible. Secondly, the UMI used in gRNA sequencing allows for discrimination between sequencing errors and mutations. GiRAFR analyses can thus identify sources of gRNA mutations. Inspecting the mutation pattern along the gRNA sequence revealed that most mutations are spread across at the gRNA spacer region, corresponding to the synthesized oligonucleotide fragment. This suggests that these mutations result from inaccurate oligonucleotide synthesis, and highlight the potential tradeoff between cheaper synthesis at lower accuracy, and the downstream information loss due to inaccurate oligonucleotide synthesis. Notably, the relatively high cost per cell in single cell CRISPR screens renders this a more pressing matter than in regular CRISPR screens, where mutant gRNAs are typically disregarded as they have a marginal downstream opportunity cost. Thirdly, GiRAFR-identified mutations may allow for an analysis of the impact of mutations on gRNA functioning. Indeed, we demonstrate that mutations in both the spacer and the scaffold compromise gRNA activity, as both are associated with a reduced ability to downregulate target gene expression. Although beyond the scope of the current study, more in-depth analysis of high-throughput datasets may enable a more fine-grained appraisal of which mutation types and locations are tolerable or damaging. Finally, we demonstrate that both the number of gRNA UMIs identified per cell as well as the gRNA library sequencing depth affect the ability to detect gRNAs mutations and to reliably assign gRNAs to cells. Our data suggest that a high sequencing saturation facilitates identifying gRNA mutations. But more importantly, the average number of gRNA UMIs differed dramatically between experiments. This may either be due to low gRNA expression or to low detection rates, but can have a detrimental impact on the accuracy to assign gRNAs to cells, to discriminate ambient from endogenously expressed gRNAs and singlets from multiplets. Together, we believe that these notions support the need for accurate gRNA calling as implemented in GiRAFR, and that this novel software tool will be of benefit to better assess the outcome of emergent single cell CRISPR screens.

## Methods and algorithm

### gRNA design

gRNAs were designed using Benchling [Biology Software, 2018], retrieved from https://benchling.com. Final gRNA sequences contained homology arms (5’-end: TGGAAAGGACGAAACACCG, 3’-end: GTTTTAGAGCTAGAAATAGCAAGTTAAAATAAGGC) to allow for cloning into the CROPseq-Guide-Puro vector (addgene plasmid #86708 from Christoph Bock). The gRNA library was synthesized and ordered as an oligo pool through CustomArray (GenScript).

### gRNA cloning

Cloning of the pooled gRNA library, and all consecutive steps until library preparation, were performed according to the CRISPR droplet sequencing (CROP-seq) protocol as detailed (3). Briefly, the vector was digested with 20 units of *BsmBI* (NEB cat. no. R0580L), and the backbone was purified using SNAP UV-Free Gel Purification kit (Fisher Scientific cat no. 45-0105). The gRNA library pool was cloned into the resulting backbone fragment with NEBuilder HiFi DNA Assembly Master Mix (NEB cat no. E2621S) and transformed into Endura Electrocompetent cells (Lucigen cat. No. 60242). Plasmids were isolated and purified using the QIAprep Spin Miniprep Kit (Qiagen, cat. no 27104).

### Lentivirus production

Lentivirus was produced by transfecting HEK293T cells using lipofectamine 3000 (ThermoFisher cat. no. L3000015) with pMDLg/pRRE, pRSV-Rev and pMD2.G (Addgene #12251, #12253, #12259) and the generated CROPseq-Guide-Puro gRNA library. For lentiviral titration, puromycin (Cayman Chemicals, cat no. 13884) resistant colonies were counted through Crystal Violet (Sigma Aldrich, cat no. 61135) staining.

### Transduction of A549-TetOn-Cas9 cells

A549 cells with a doxycycline inducible Cas9 construct were created by transducing A549 cells with lentivirus encapsulating the pCW-Cas9-Blast vector (Addgene, #83481), followed by selection with 20 μg/mL blasticidin (InvivoGen, cat no. 38220000). The A549-TetOn-Cas9-cells were transduced with the gRNA library pool lentivirus at a multiplicity of infection of 0.3. Positively transduced cells were selected for with puromycin (100 μg/mL). Following selection, Cas9 expression was initiated by doxycycline (VWR, J60579.14) addition (5 μg/mL). After two days of induction, cells were grown for another 5 days, collected and processed following the 10x Genomics demonstrated protocol for the preparation of single cell suspensions. Cells were grown in DMEM high glucose (Thermo Fisher, 41965062) at 37 ̊C in 5% CO2 and passaged every 2 days with Trypsin-EDTA (Thermo Fisher, 25200056).

### CROPseq library preparation

~5,000 cells were processed with Chromium Next GEM Single Cell 3’ GEM, Library & Gel Bead v3.1 kit (10X Genomics, 1000128) following the manufacturer’s instructions. The final library was analyzed using bioanalyzer high sensitivity DNA analysis kit (Agilent, 5067-4626). Final library was sequenced at NovaSeq 6000 (R1:28 and R2:91).

### gRNA enrichment for library preparation

10% of cDNA was used for gRNA enrichment and creation of a specific gRNA library. For higher resolution and better cell assignments, gRNA sequences were amplified with Hifi HotStart ReadyMix (Roche, 7958935001) and gRNA cassette specific primers (5’-CAAGCAGAAGACGGCATACGAGATTAAGGCGAGTGACTGGAGTTCAGACGTGTGCTCTTCCGATCTTCTTGTGGAAAGGACGAAACACCG-3’ and 5’-AATGATACGGCGACCACCGAGATCTACACTCTTTCCCTACACGACGCTCTTCCGATCT-3’) before the last indexing PCR. Final gRNA library was sequenced together with the transcriptome library.

#### CROPseq data process

We performed CRISPR droplet sequencing (CROP-seq) using 4 different gRNAs for each of 25 target genes, and with 20 non-targeting guides as controls. Gene expression library: 454,191,138 reads, CRISPR library: 25,289,660. Alignment of gene expression library, detection of spacer and cell assignments were performed by Cell Ranger (version 3.1.0) feature barcoding analysis. The count matrix of cells with a single spacer was then extracted and normalized using LogNormalize function (R version 4.2.0 (15) and R package Seurat, version 4.1.0 (16)) with a scale factor 10,000 for further analysis.

#### GiRAFR

##### Generation of custom reference genome

A custom reference genome was built using Cell Ranger mkref (build note can be found in user manual). It was composed of the human reference genome GRCh38, supplemented with an artificial chromosome for each gRNA, containing the gRNA-specific sequence as well as upstream and downstream sequences relevant for the underlying method. Specifically, for data using 10X protocol, we used 530bp from the plasmid containing the gRNA cassette. For data originating from Dropseq, we included the requisite Dropseq-tools index in the custom reference genome file.

##### Mapping of reads to the custom reference genome and filtering of mapped reads

Next, reads from the targeted sequencing library and the transcriptome sequencing library were aligned to this custom reference genome by STAR as implemented in Cell Ranger or Dropseq-tools. For 10x sequencing, Cell Ranger performed correction on cell barcode and UMI with a maximal 1 hamming distance. All subsequent analyses were limited to reads having both a valid cell barcode and a valid UMI (CB and UB tag respectively for 10x and XC and XM tag for Dropseq). After removal of the secondary alignments, reads mapping to the artificial gRNA chromosomes were collected in a BAM file.

This bam file was filtered by the predefined cell barcodes list, containing cell barcodes that remain after removal of background noise. In most cases, this list came from alignments of gene expression library. To solve the cell barcodes discrepancy between gene expression library and captured CRISPR library by 10X feature barcoding technology, unmatched cell barcodes were compared and corrected with lookup table provided by Cell Ranger.

##### gRNA consensus sequence calling

Part of the reads in the gRNA filtered BAM are associated with an identical UMI and cell barcode. These are assumed to originate from the same cDNA molecule. Because of errors in library preparation and/or sequencing, the associated nucleotide sequence in these reads is not necessarily identical. We therefore defined a consensus sequence for each UMI-cell barcode combination. Specifically, Pysam (9) (Python package version 0.15.3 (17)) was used to extract the set of reads for UMI-cell barcode combination from the gRNA filtered BAM file. Firstly, the mapped gRNAs of this set of reads were inspected (GN tag for cell ranger and gn for dropseq-tools bam file). In most of cases, they align uniquely to one gRNA. In case of multiple gRNA annotations for the same UMI-cell barcode combination, the most reads supported gRNA was kept. In case of a tie, no gRNA can be confidently mapped. Those UMIs were labeled as ‘multiple genes’ and kept out from gRNA calling. From the subset of reads that unanimous mapped, the most common sequence was defined as the consensus sequence for this UMI-cell barcode combination. When multiple different reads have identical counts, we randomly selected the consensus sequence from this set. Consensus sequence of the entire set of reads was given when no confident gRNA mapped.

##### Filtering of gRNA consensus sequences

In theory, consensus reads from UMI-cell barcode combinations that were constructed using multiple reads can be considered as of higher confidence, because this excludes sequencing or other errors. In GiRAFR, we provide the option to filter consensus sequences according to the number of reads supporting them (default: > 1). Cell barcodes, UMIs and corresponding consensus sequences were written into “consensus.sequence.gRNA.txt” file. Alignments with consensus sequenced were also saved into a “consensus.bam” file which is compatible to other analysis tools.

##### Identification of mutations in the gRNA consensus

Analyzing the alignment information is a fast and accurate way identifying mutations in short sequences such as in the oligonucleotides used to construct gRNA libraries. With the designed gRNA and cassette sequence annotation, we detect aberrations per base wise. From the CIGAR strings in BAM alignments files, GiRAFR keeps all insertions (I), deletions (D), skipped regions (N), soft clipping (S) and compare mapped bases to references (M). GiRAFR can also annotate the structure affected by the mutation using oligo structure annotation provided by the user and detailed mutation in a CIGAR similar way (details in supplementary). Perfect match without any mismatch will be defined as intact UMI. Variant guides are also named with number suffix.

##### Assign gRNAs to cells

Ambient gRNAs molecules can be captured in droplets and to erroneously assigned to cells. To avoid this, a threshold defining the minimal number of UMIs supporting the assignment of a gRNA to a cell is defined. Two modes are available. Users can define a fixed threshold (for example 3 UMIs) which will be applied to all gRNAs. Alternatively, automatic dynamics detection of UMI thresholds can be employed. It will fit a two-components mixture gaussian model to model gRNA UMIs and background noise (“ambient”) UMIs. It sets the UMI threshold for a cell to have gRNA as the minimum UMI of the gRNA UMI counts distribution. Intact gRNA and their respective variants are counted together in fitting this distribution. This method is inspired and adopted from Cell Ranger. Only gRNAs in a cell with more than the minimum UMI threshold will be assigned to that cell. The resulting output consists of a matrix containing the UMI counts per guide (including mutated guide) and per cell.

##### Detect CRISPR cas9 editing effect

Alignments from expression library are first split into sub bam files to accelerate detection speeds by SAMtools (version 1.9)(18). Each sub bam file contains the alignments that mapped over the detection window of each gRNA. The detection window is short and symmetric around the Cas9 nuclease cutsite on the genome, which is 3-4nt upstream of the PAM site. We used a 51 bp detection window. Tne FeatureCounts (v2.0.1) (19) tool was used to annotate mapped reads to genes. Only alignments which are assigned and mapped to target genes are considered valid.

Similarly, to deduplicate reads with identical UMI and cell barcode, we construct a consensus sequence. To address that reads do not align to the same starts within the detection window, we take unions of sequenced reads as the consensus sequence for each UMI cell barcode combination. By analyzing mapped positions of consensus sequences, if deletions happen across cutsite, then they are detected as editing effects. Insertions or deletion elsewhere and mismatches are not considered as CRISPR-Cas9 induced are be reported in the final outputs. gRNA assignment of the cell from previous step is also reported.

## List of abbreviations

KO: Knockout
scRNA-seq: Single-cell RNA-seq
GiRAFR: Guide RNA Aberrations and Functionality Revealed
CROP: CRISPR droplet sequencing
sgRNA: Single guide RNA
UMI: Unique molecular identifiers
TSO: Template switch oligo
PAM: Protospacer adjacent motif

## Availability of data and materials

GiRAFR is available as an open-source Python script at our GitHub Repository (github.com/FunctionalEpigeneticsLab/GiRAFR).

CROP-seq data for this study has been deposited in the NCBI’s Gene Expression Omnibus (GEO) database under the accession number GSE216040.

## Authors’ contributions

Q.Y. wrote the software as well as ran the analyses detailed in this manuscript. Q.Y. and B.T. conceived the project. Q.Y., B.T. wrote the manuscript with contributions from P.V.M. E.G performed experiments for the In House proof of concept screen. P.V.M. provided feedback on the interpretation of the results. All authors read and approved the final manuscript.

## Acknowledgements

We are grateful to all members of the Thienpont lab for the valuable discussions. We thank all authors of the data used in this study for their previous work and making their results available.

## Supplementary Information

### CIGAR-like string

Digit numbers represents exact matches, and nucleotides followed are mutated bases. 0 represents no nucleotide.

Digit numbers followed by insertions (I), deletions (D) and soft clippings (S) show the number of nucleotides of those events. Hard clippings (H) are not included. The major difference between this string and CIGAR-string is it replaces matches (M) into mismatches and encode detailed mutated nucleotides [ATGC] into the string.

### Mutation structure annotation

Annotations begin with oligo structures such as gRNA which are consistent with user input oligo pool plasmid structure.gtf. Then each mutation annotation follows oligo structure with semicolon as separator. Comma separates individual mutation event. Digit numbers represents the distance to the beginning of the structure. Nucleotides followed are mutated bases. 0 represents no nucleotide. Digit numbers in bracket followed insertions (I), deletions (D) and soft clippings (S) represent the number of nucleotides of those events.

### Supplementary Figures

**Figure S1.**
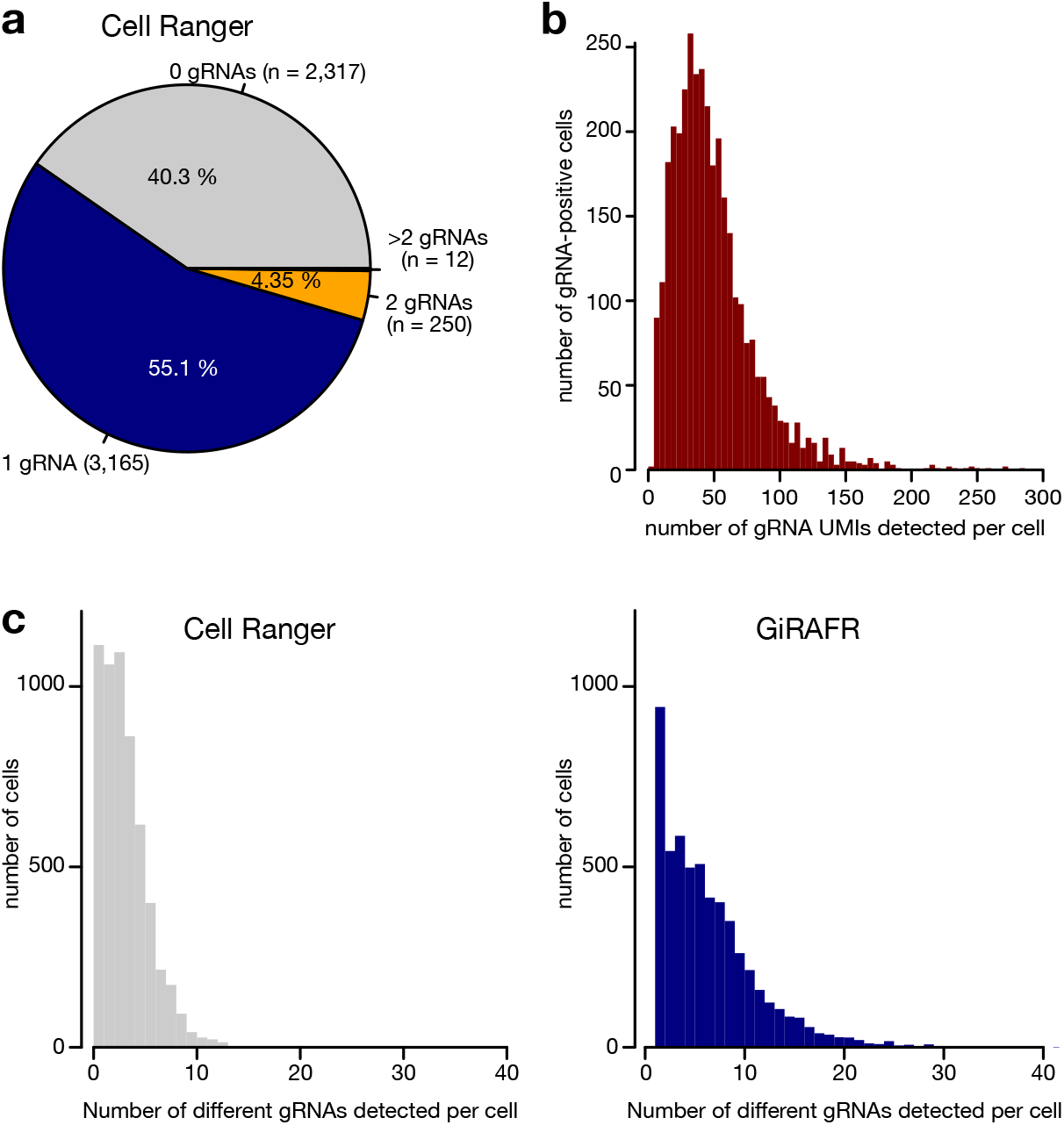
**a**. Pie chart showing the number of cells with 0, 1, 2 or more than 2 gRNAs assigned to them by Cell Ranger. **b**. Histogram showing the number of different gRNA UMIs detected in gRNA-positive cells by Cell Ranger. **c**. Histograms showing the number of different gRNAs detected by Cell Ranger and GiRAFR. Note that GiRAFR discards all cells in which no gRNA is detected. For Cell Ranger, only gRNAs matching the designed sequence are considered, while for GiRAFR, both intact and mutant gRNAs may be counted.

**Figure S2:**
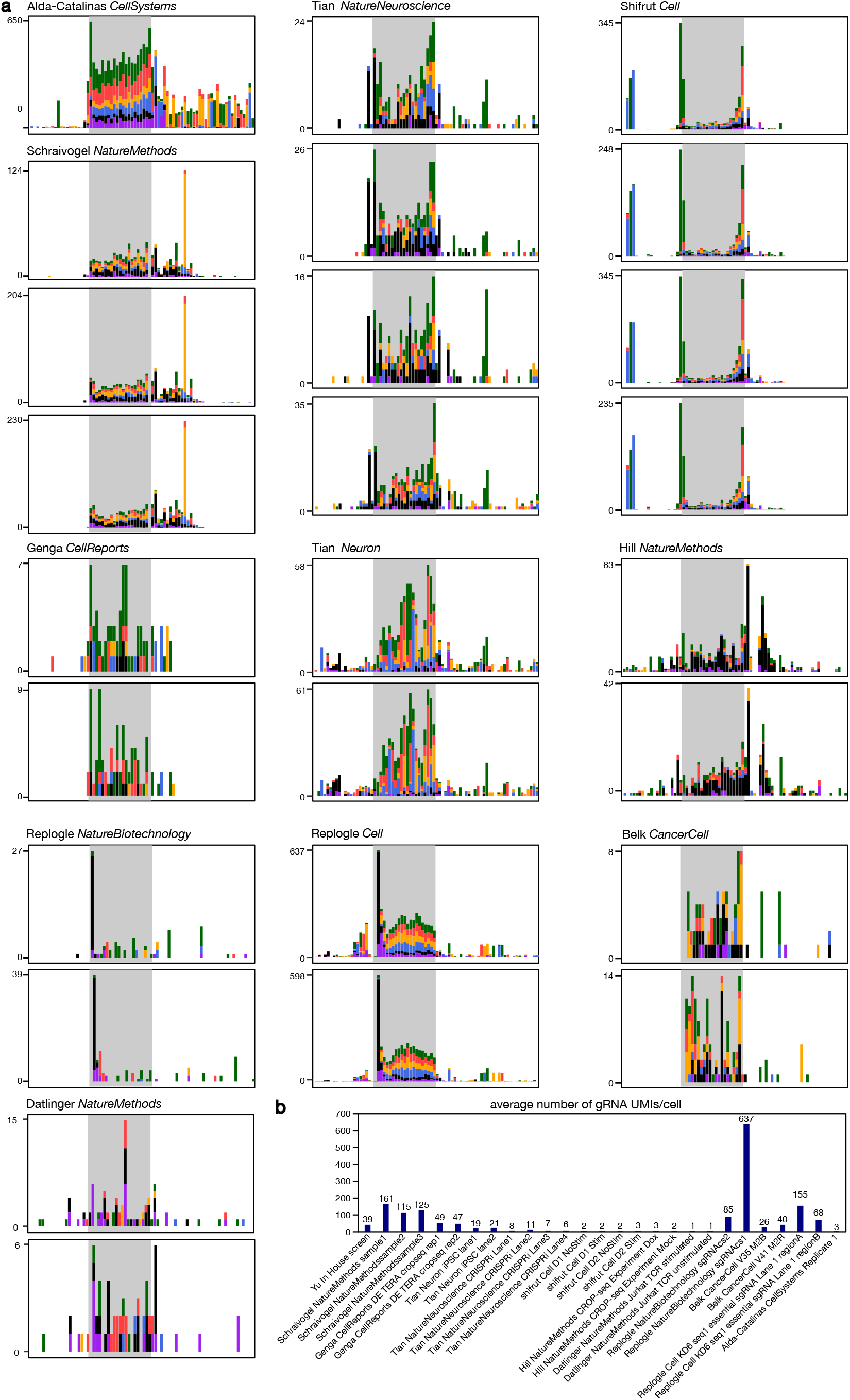
**a**. Mutation frequency of gRNAs associated with cells after the Gaussian Mixture model filtering. Shown are all gRNA mutation patterns detected in publicly available datasets. Note that some gRNA libraries encompass only a fraction of the U6 promoter shown. The aggregate of these experiments per study is shown in Figure 2e. Striped boxes indicate the position of template switching oligonucleotides, where mutations on position −2, −1 and 0 before the start of the spacer were also removed. Such mutations are likely produced by the terminal nucleotidyl transferase activity of the reverse transcriptase, and hence not present in the gRNA. **b**. the average number of gRNA UMIs per cell, as determined by GiRAFR.

**Figure S3:**
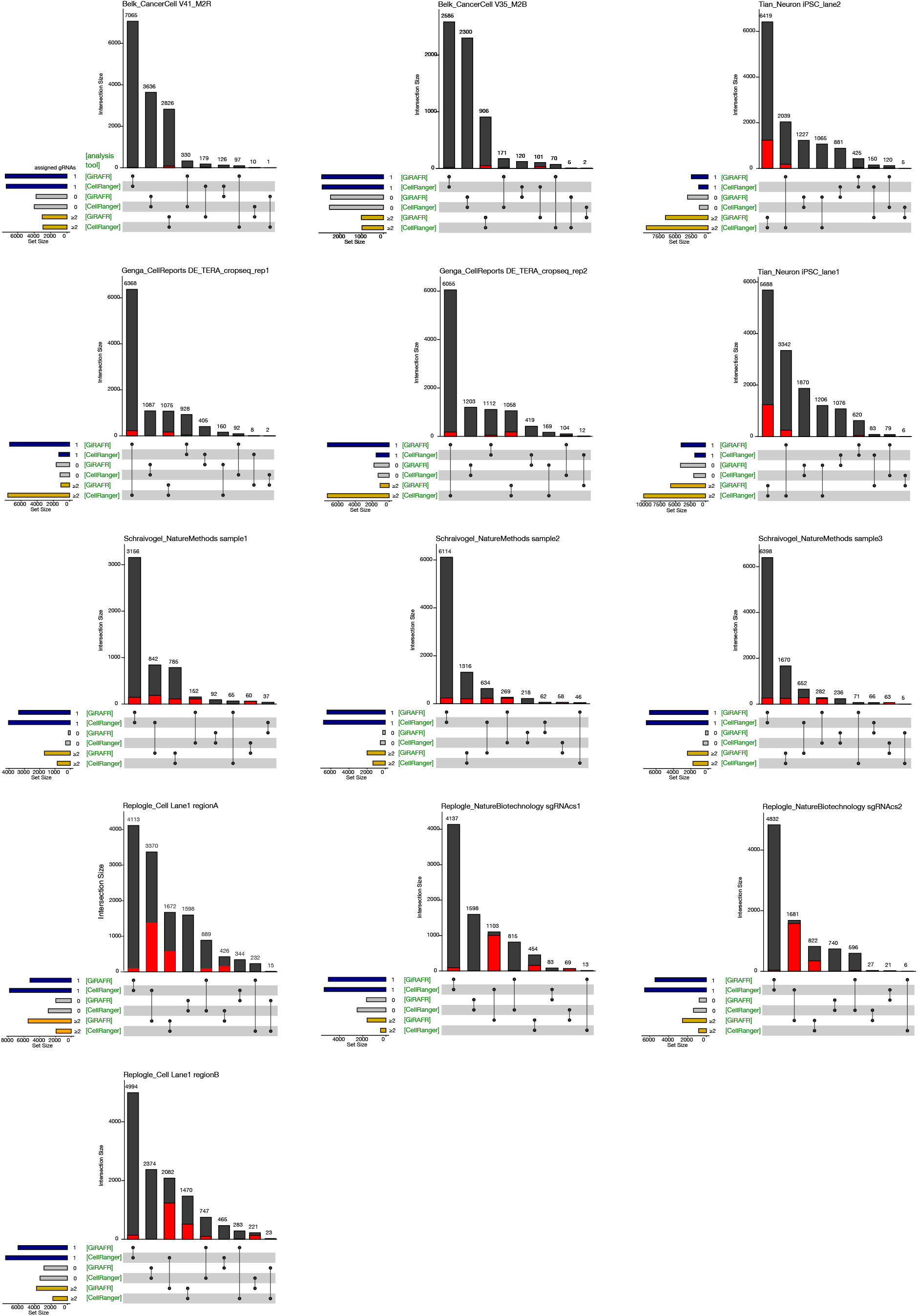
Comparison of gRNA assignment to cells between Cell Ranger and GiRAFR (20). The red box indicates cells containing 1 or more mutant gRNAs. Shown are data for each of the 13 experiments for which on average over 16 unique gRNA molecules were recovered per cell, suggesting that over 95 % of cells are correctly assigned.

**Table S1.**
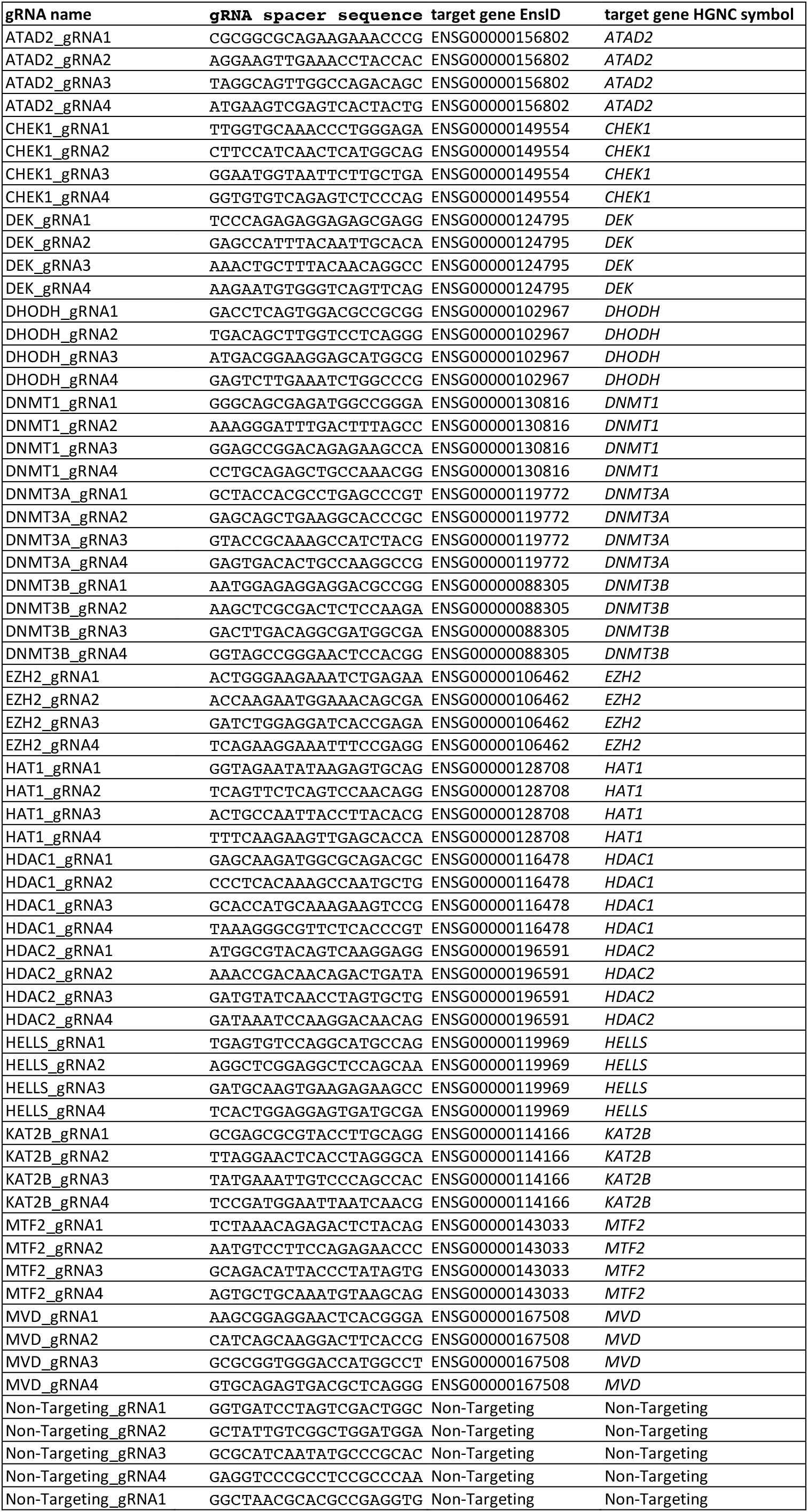
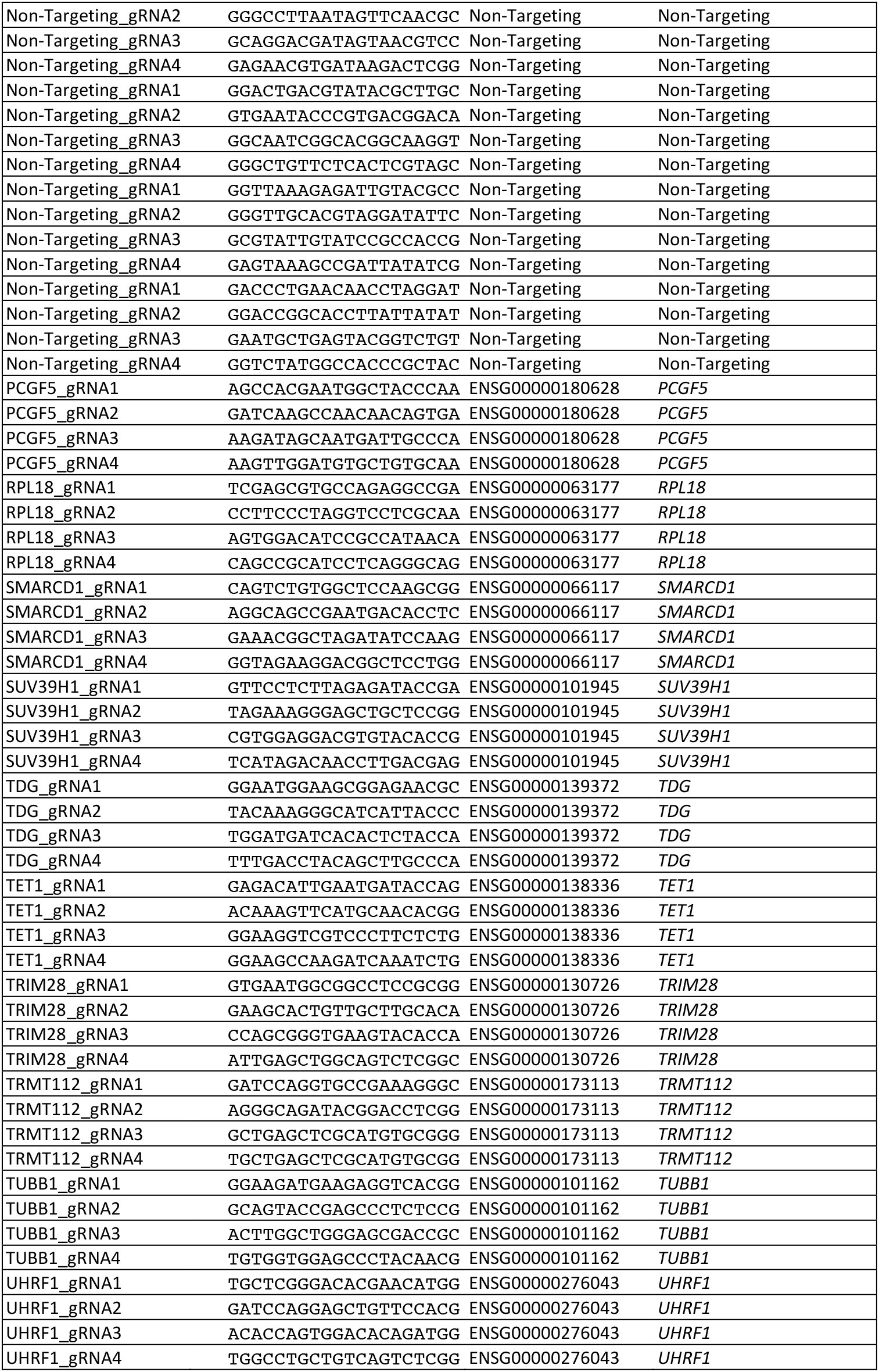

**Table S1.**
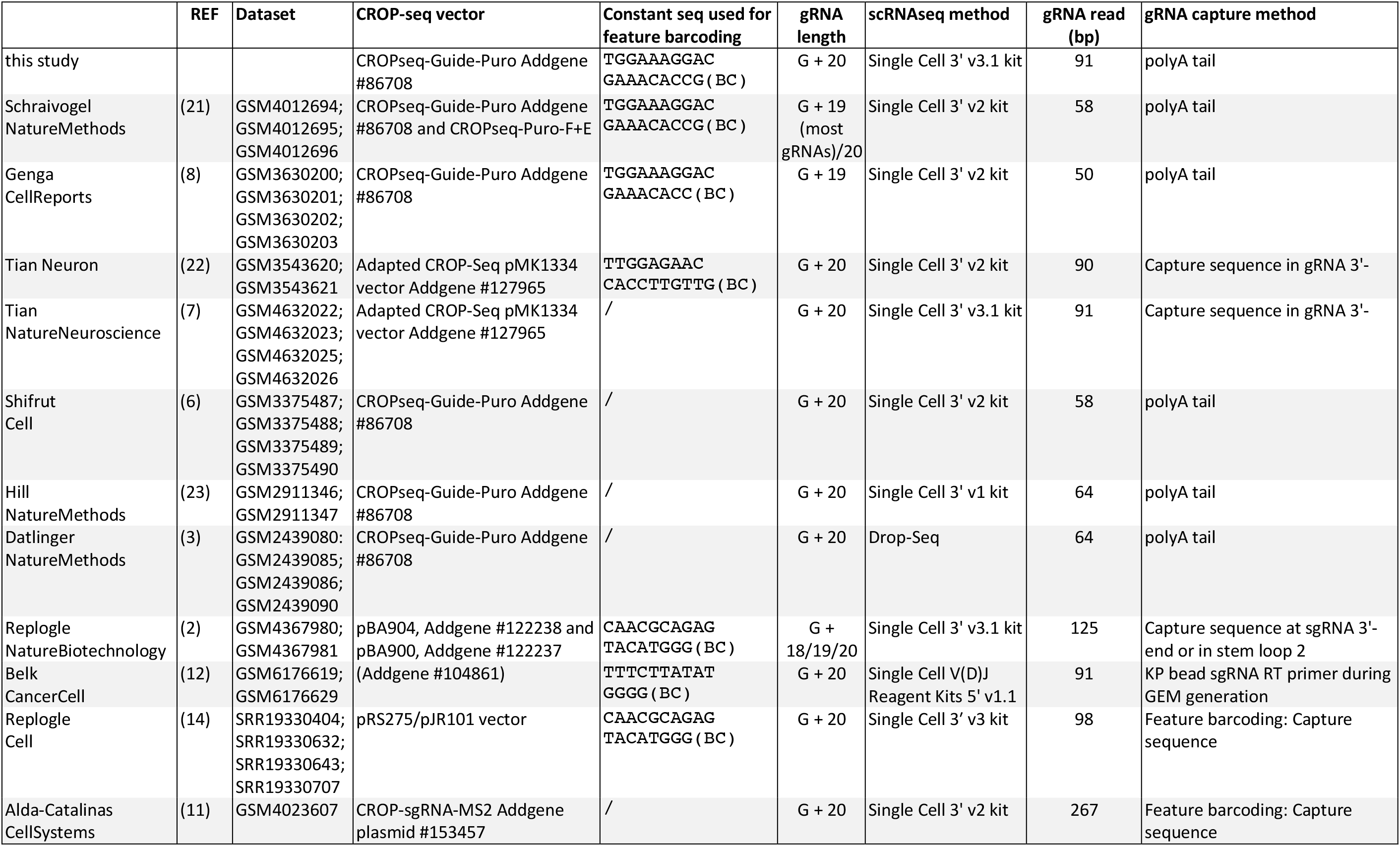

